# Metacontam: A Negative Control-Free Decontamination Method for Metagenomic Analysis

**DOI:** 10.64898/2026.04.26.720876

**Authors:** Junwoo Jo, Hyeyoung Lee, Jae Woo Baek, Suyeon Lee, Vikas Singh, Adil Mardinoglu, Saeed Shoaie, Jinwook choi, Sunjae Lee

## Abstract

Shotgun metagenomic sequencing enables high-resolution profiling of host-associated microbial communities. However, contaminant DNA can substantially distort biological interpretations, especially in low-biomass samples. Here, we introduce Metacontam, a control-free method for species-level decontamination of shotgun metagenomic data. Metacontam integrates blacklist-guided community detection within a species correlation network with average nucleotide identity (ANI) to identify contaminants arising from shared sources. Across diverse low-biomass and mixed-biomass datasets, Metacontam outperformed existing approaches, improving the detection of low-abundance and low-prevalence contaminants while retaining biologically plausible taxa. It also reduces kit-specific biases in skin metagenomes and improves downstream analyses of tissue microbiome data. Together, these results demonstrate that Metacontam enables accurate identification of contaminant taxa across diverse metagenomic datasets, even in the absence of negative controls.

## Introduction

Over the past decades, whole-genome shotgun (WGS) sequencing has advanced our understanding of human-associated microbiota, including their composition and functional potential in health and disease. Unlike targeted amplicon sequencing, WGS metagenomics has enabled in-depth taxonomic profiling that provides high-resolution characterization of microbial community composition (e.g., species/strain-level profiling) and functions (e.g., orthologous genes)¹.

However, in low-biomass microbiomes, including the skin and nasal cavity, as well as in tissue biopsy samples, contamination from exogenous microbial DNA poses a major challenge for accurate microbial taxonomic profiling in shotgun metagenomic studies^2–6^. Unlike high-biomass samples such as the gut or oral cavity, where abundant endogenous microbial DNA can mitigate the influence of contaminants, low-biomass samples are highly susceptible to contaminant signals introduced through reagents or laboratory environments^7^. If not properly identified and removed, such contamination can distort microbial community profiles and lead to inaccurate biological interpretation^8–11^.

Despite efforts to address contamination in low-biomass metagenomic studies, existing computational approaches for contaminant removal suffer from practical limitations^4,12–15^. For example, Decontam, a recently developed tool, provides two approaches for identifying contaminant DNA: prevalence-based and frequency-based methods. However, Decontam, like most available methods, depends on negative controls or DNA concentration measurements. Generating negative controls for every sample at each stage of the experimental workflow is costly and impractical. Moreover, in many studies, negative controls are either not generated or not publicly available.

Squeegee is the only tool capable of identifying contaminant taxa without negative controls^16^. It identifies the contaminant species observed across compositionally distinct sample types that show high prevalence and broad genome coverage. However, its performance relies on the availability of taxonomically distinct sample types and has not been validated for datasets composed solely of low-biomass samples, which is a common scenario in microbiome studies. When negative controls are unavailable, a commonly used alternative is a microbial blacklist, a curated collection of taxa detected as contaminants in metagenomic studies^4,17–19^. However, blacklist-based filtering fails to identify study-specific contaminants^4^.

Another complementary approach is to exploit interspecies correlation structures for the identification of contaminant species. Contaminant species are expected to co-vary across samples and often exhibit strong correlations, thereby providing an initial signal of contamination^7^. However, each species can correlate with many others, yielding a dense web of edges. Therefore, a single correlation is insufficient to classify a species as a contaminant, and correlations involving low-abundance species are often unreliable^20,21^. Moreover, it is difficult to generalize the correlation cutoffs across different datasets. Therefore, we reasoned that a network-based view of species associations could simplify this complexity and reveal contaminant-associated clusters.

Real-world biological networks, including microbiomes, are highly structured rather than random and exhibit communities that influence system behavior^22–25^. To characterize the structural properties of microbial communities, we applied community detection, a network-based approach that identifies groups of nodes that are densely connected to one another and sparsely connected to the rest. These groups are commonly referred to as communities (also termed clusters or modules) ^26–28^. We hypothesized that contaminant species would co-cluster into distinct community within a species association network.

Here, we introduce Metacontam, a decontamination method that leverages microbial community structure and species-level genomic similarity to identify contaminant taxa without negative controls. Metacontam first performs community detection on a species correlation network guided by known blacklist taxa to identify a contaminant community. The identified community is then refined using ANI, as contaminants originating from a common external source tend to exhibit higher genomic similarity across samples than genuine taxa. We validated Metacontam across diverse datasets containing low-biomass samples, including breast milk, placenta, skin swabs, nasal swabs, and tissue biopsies. Across all datasets, Metacontam outperformed Decontam and Squeegee, effectively removing low-abundance contaminant species while retaining genuine taxa. Through computational and experimental simulations, we further confirmed the robustness and core assumptions of Metacontam. Taken together, Metacontam will advance shotgun metagenomic studies of low-biomass samples, enabling broader application even in the absence of negative controls.

## Results

### Development of a network- and ANI-based decontamination approach

Recent advances in high-resolution shotgun metagenomics have enabled low-biomass microbiome analysis. However, microbial communities should be understood in the context of their ecological interactions within specific spatial niches^59^. While bulk metagenomic analyses provide a general overview of community composition, they offer only a rough resolution of these interactions. To fully capture genuine ecological interactions, one must examine the micro-niches in which these communities are assembled - making low-biomass microbiome studies an unavoidable choice.

Currently, most decontamination tools rely on negative controls or simple prevalence metrics and fail to effectively identify contaminants in low-biomass samples^4,16^. To address this, we assumed two key aspects of contaminant taxa: (1) contaminant species originating from a shared external source tend to co-occur and form positively correlated cluster within a species association network, and (2) contaminants from a common source exhibit higher genomic similarity across samples, measured by average nucleotide identity (ANI), whereas genuine species derived from diverse host environments show greater genomic variation. Based on these assumptions, we devised Metacontam, a decontamination pipeline integrating two modules described below.

### Community Detection Module

Based on our assumptions, we devised a contaminant community detection approach using the Louvain algorithm guided by a curated blacklist of known contaminant species, which enables effective selection of contaminant communities (**Fig. 1a**). First, we performed species-resolution taxonomic profiling using Kraken2^52^ and Bracken^53^, which classify metagenomic reads and estimate species-level abundances, respectively, and constructed a species correlation network (Pearson correlation on CLR-normalized profiles; see **Methods** for network construction details). Second, we conducted a two-phase Louvain procedure: in the seeding phase, all blacklist species were unconditionally grouped into a single seed community regardless of modularity gain, with intra-community edges converted to self-loops and inter-community edges aggregated as weighted edges. The subsequent Louvain phase then proceeded identically to the original algorithm, iteratively recruiting correlated taxa into the seed community until no further modularity gain was achievable. The resulting community containing blacklist species was defined as the contaminant community.

**Fig. 1.**
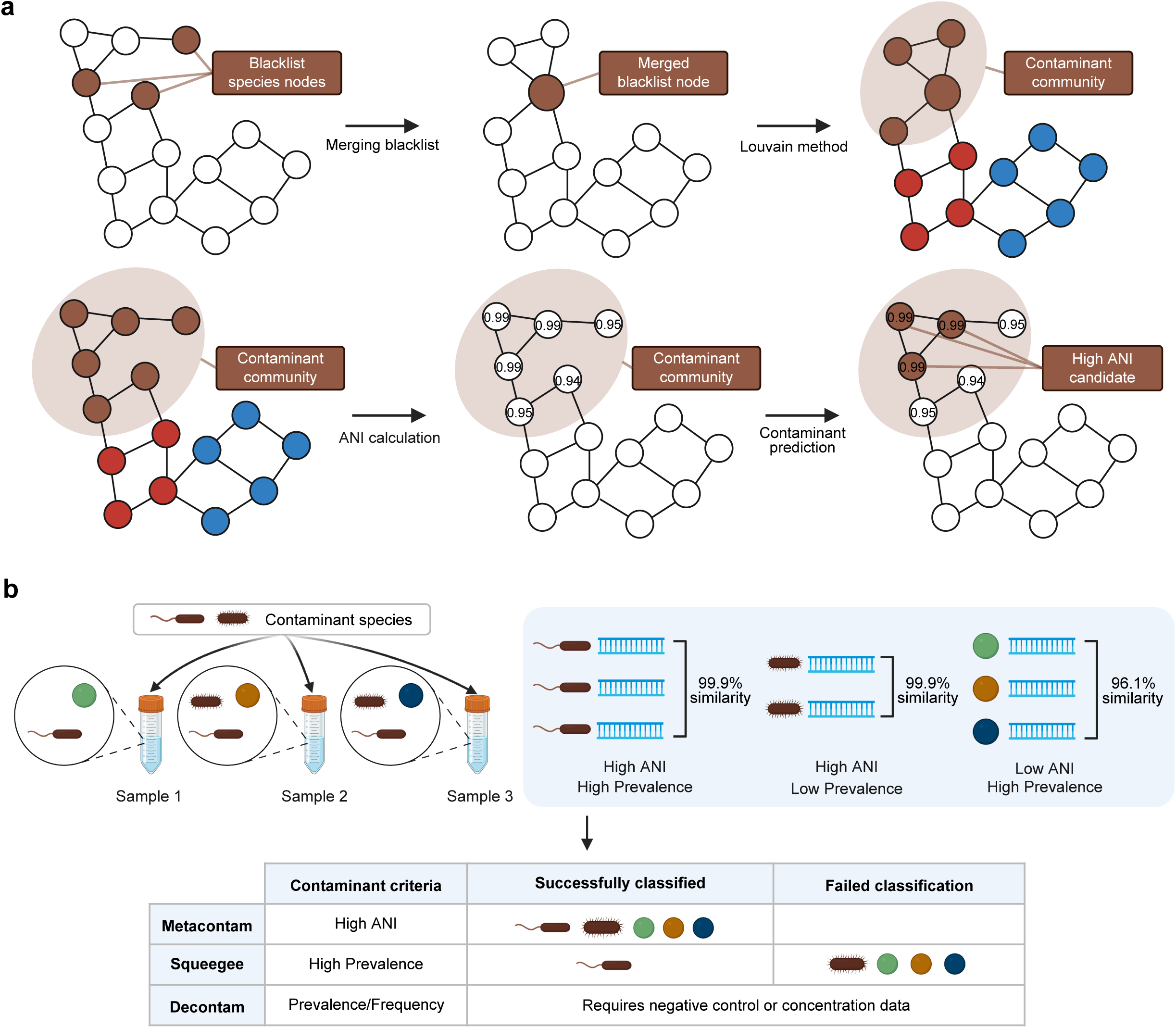
Conceptual framework and workflow of Metacontam. **a**, Workflow of Metacontam. Blacklist species are first identified in the species association network and collapsed into a single seed node. Louvain community detection is then applied to identify the contaminant community. Within this community, ANI is calculated for each species across samples, and species with a mean ANI above the predefined ANI threshold are flagged as final contaminants. The rule used to define the ANI threshold is described in the **Methods**. **b**, Schematic illustration of the ANI-based rationale used in Metacontam in the absence of negative controls, together with a comparison table summarizing the criteria used by Metacontam and existing methods for contaminant identification.

### ANI-based Species Tracking Module

Next, based on our assumption that contaminants from a shared external source exhibit higher genomic similarity, we devised an ANI-based species tracking module. For species within the identified contaminant community, we quantified genomic similarity across samples, and species with high mean ANI were flagged as candidate contaminants (**Fig. 1a**; see **Methods** for threshold criteria). For datasets with more than 45 samples, 1,000 sample pairs were randomly selected using Mash^56^ distance-stratified sampling to reduce computational cost. **Figure 1b** illustrates the rationale for ANI-based contaminant identification in the absence of negative controls and highlights the limitations of prevalence-based approaches such as Squeegee, which may miss lower-prevalence contaminants and misclassify highly prevalent bona fide species as contaminants. As ANI needs alignment computations, we also provided runtime of Metacontam (**Supplementary Table 2**, approximately 1.4 hour for 10 samples). In addition, we also provided the metadata of shotgun metagenomics samples where we applied Metacontam in this paper (**Supplementary Table 3**).

### Evaluation of Metacontam for contaminant community detection in the maternal–infant dataset

We evaluated Metacontam on a benchmarking dataset from Liu et al.¹L, which is particularly well-suited for this analysis as it encompasses diverse sample types (e.g., vaginal, fecal, oral, placental, and breast milk), covers both low- and high-biomass microbiomes, and includes 31 annotated ground-truth (GT) contaminant species. Using the blacklist seed (**Supplementary Table 4**), we applied the Louvain method and identified community 5 (C5) as the candidate contaminant community. Of the 31 GT species, 26 were present in the network (83.9%), and all 26 were assigned to C5 (**Fig. 2a**). We further examined the robustness of Metacontam to blacklist size and found that as few as 5 blacklisted species were sufficient to recover all 26 GT species into a contaminant community (**Supplementary Fig. 1a**), demonstrating that the blacklist-guided Louvain algorithm effectively captures contaminant species even with a small blacklist.

**Fig. 2.**
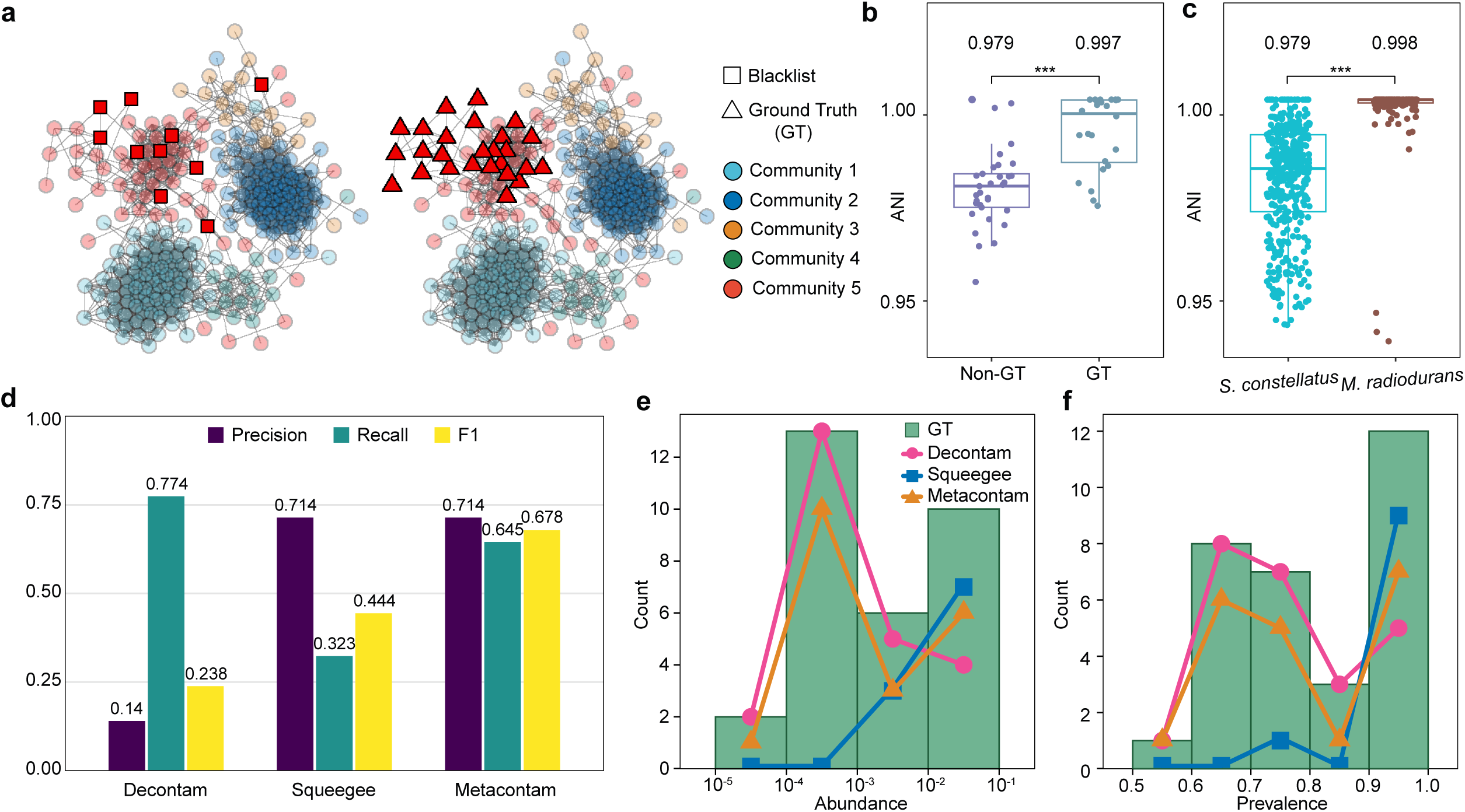
Metacontam improves species-level contaminant detection using community structure and ANI. **a**, Microbial association networks for the mother–infant dataset. Edges denote positive Pearson correlations (r ≥ 0.45). Nodes are colored by community (only communities with ≥10 nodes are shown). Panels share the same layout: left, blacklist taxa (squares); right, ground truth (GT) contaminant taxa (triangles). All other taxa are shown as semitransparent circles. **b**, ANI distributions for GT contaminants and non-GT taxa within community 5 (C5). Each point indicates the per-species mean ANI across samples, and the values above the boxes indicate the medians. One-sided Wilcoxon rank-sum test. **c**, ANI distributions for *Streptococcus constellatus* and *Methylobacterium radiodurans*. Each point denotes the ANI for the sample pairs, and the values above the boxes indicate the means. One-sided Wilcoxon rank-sum test. * p ≤ 0.05; ** p ≤ 0.01; *** p ≤ 0.001; ns, p > 0.05. The exact P-values are provided in **Supplementary Table 1**. **d**, Benchmark comparing Decontam, Squeegee, and Metacontam. The bars show the precision, recall, and F1 scores for each method. **e,f**, Distribution of GT contaminant species and true positives identified by each method based on abundance (**e**) and prevalence (**f**). The x axis in **e** is shown on a log10 scale.

Within C5, we assessed genomic similarity across samples using ANI. As expected, GT species exhibited significantly higher mean ANI than non-GT species (**Fig. 2b**), confirming our assumption that contaminant species originating from a shared external source show reduced genomic diversity across samples. For example, *Methylobacterium radiodurans* (GT contaminant; blacklist) displayed markedly higher ANI compared to *Streptococcus constellatus* (oral commensal; non-contaminant) (**Fig. 2c**).

### Benchmarking against Decontam and Squeegee in the maternal-infant dataset

Next, we benchmarked Metacontam against Decontam and Squeegee using a maternal-infant dataset. Metacontam achieved precision/recall/F1 = 0.714 (20/28 species), 0.645 (20/31 species), and 0.678. In comparison, Decontam achieved precision/recall/F1 = 0.140 (24/171 species), 0.774 (24/31 species), and 0.238, while Squeegee achieved precision/recall/F1 = 0.714 (10/14 species), 0.323 (10/31 species), and 0.444. Overall, Metacontam outperformed both methods in terms of F1 score (**Fig. 2d**). Notably, Decontam achieved higher recall but at the cost of substantially lower precision, whereas Squeegee achieved high precision but a much lower recall. In contrast, Metacontam achieved higher precision than Decontam and higher recall than Squeegee, demonstrating a more balanced precision-recall trade-off and resulting in the highest F1 score among the three methods.

Next, we assessed whether performance improved with the incremental addition of three method components (**Supplementary Fig. 8a**): (i) blacklist-based identification (predicting contaminants using the blacklist alone), (ii) network-based community filtering (restricting predictions to taxa in the blacklist-containing community), and (iii) ANI-based refinement (further prioritizing taxa with high genomic similarity across samples). F1 increased from 0.323 to 0.553 to 0.678, with the network component primarily improving the recall and the ANI component improving precision. Together, the network and ANI components balanced recall and precision, resulting in the highest overall performance.

### Enhanced detection of low-abundance, low-prevalence contaminants

To better understand the differences in performance, we compared the abundance and prevalence distributions of true-positive (TP) taxa predicted by each method with the distribution of GT species (**Fig. 2e,f**). We observed that TPs detected by Squeegee were concentrated at higher abundance and prevalence, whereas Metacontam identified TPs across a broader range more closely resembling the distribution of GT species. Notably, Metacontam detected TP taxa spanning a wide abundance and prevalence range, including extremely low-abundance, low-prevalence taxa, such as *Methylobacterium terrae* (0.00617%; 0.541), and relatively high-abundance, high-prevalence taxa, such as *Yersinia enterocolitica* (4.65%; 0.991). In contrast, the lowest TP detected by Squeegee was *Cutibacterium acnes* (0.477%; 0.983), which was substantially more abundant and prevalent than the above *M. terrae*. While Decontam also covers a broad range, this comes at the cost of substantially reduced precision. Together, these results suggest that Metacontam more effectively captures low-abundance, low-prevalence contaminants, while maintaining a favorable precision-recall balance.

### Metacontam reduces kit-specific bias in skin metagenomes

To evaluate Metacontam in a low-biomass sample type, we reanalyzed human skin shotgun metagenomes from Shaffer et al^29^. In this dataset, skin samples were collected from the armpit, foot, forehead, and lower arm, representing distinct microenvironments within the skin dataset, and six different DNA extraction kits were compared. We selected two representative DNA extraction kits for evaluation, the MagMAX Microbiome Ultra Nucleic Acid Isolation Kit (“microbiome kit”) and the MagAttract PowerSoil DNA Isolation Kit (“standard kit”), as they exhibited the strongest kit-specific taxonomic signatures in negative controls (R² = 0.177, P = 0.001; **Fig. 3a**; **Supplementary Fig. 4**).

**Fig. 3.**
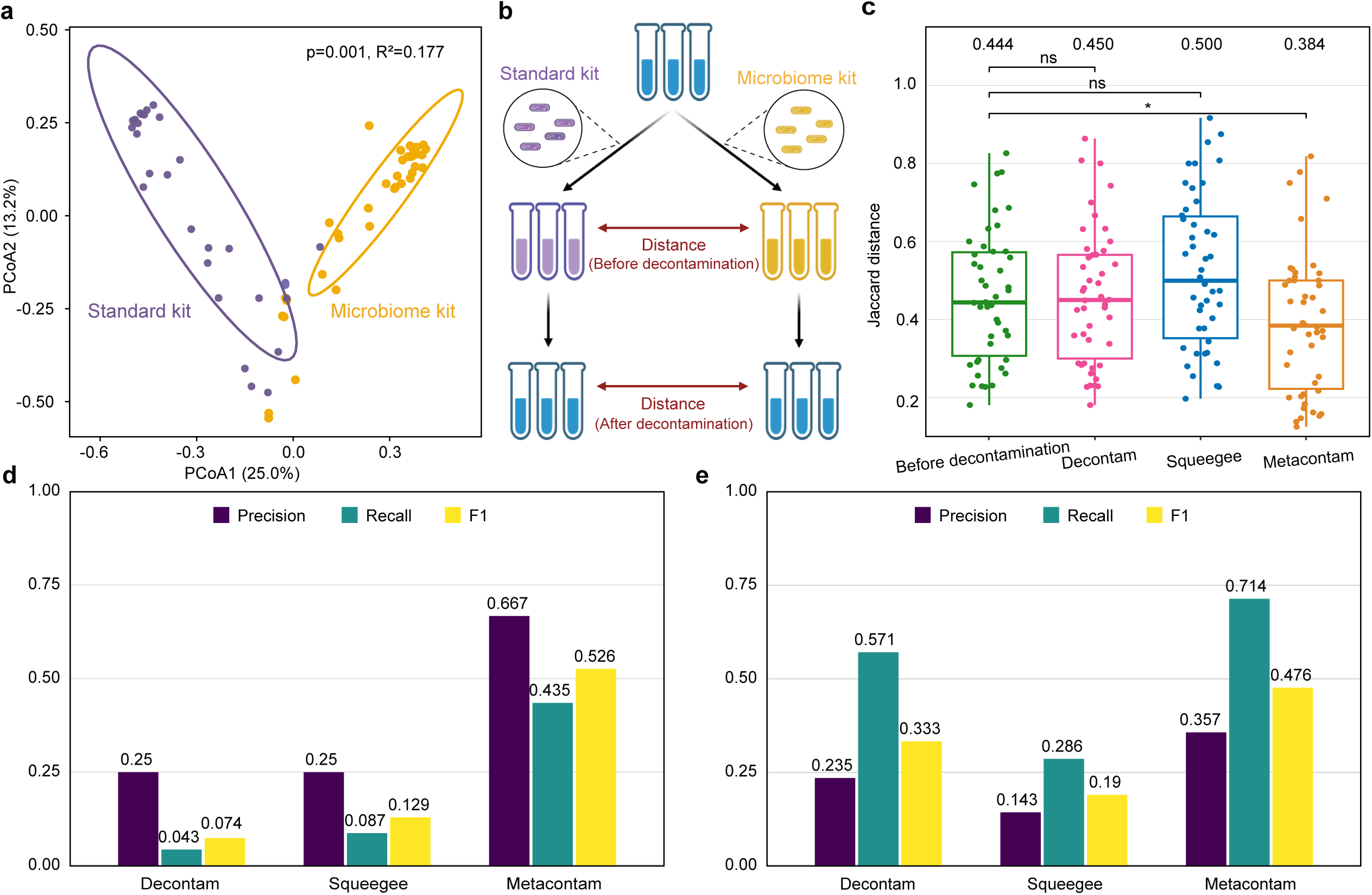
Metacontam reduces kit-specific bias in skin metagenomes. **a**, PCoA of sterile water blank samples based on Bray–Curtis dissimilarities, comparing two extraction kits: Qiagen PowerSoil (standard; *n* = 31) and MagMAX Microbiome Ultra (microbiome; *n* = 36). PERMANOVA (adonis2, 999 permutations): *p* = 0.001, *R*² = 0.177. **b**, Schematic of validation strategy. The contaminant removal methods were evaluated using matched biological replicate pairs from standard–microbiome extraction kits. Better performance is defined as a greater reduction in the distance between replicate pairs after filtering. **c**, Pairwise Jaccard distances for matched standard–microbiome replicate pairs (*n* = 46), quantifying the validation strategy shown in **b** under four strategies: Before decontamination, Decontam, Squeegee, and Metacontam (after removing predicted contaminant taxa). Taxa were retained if prevalence ≥ 0.35 and mean relative abundance ≥ 0.01. Values above the boxes indicate medians. One-sided Wilcoxon rank-sum test. p ≤ 0.05 *, p ≤ 0.01 *\*\*,* p ≤ 0.001 ***; ns, p > 0.05. The exact P-values are provided in **Supplementary Table 1**. **d,e**, Benchmark comparing Decontam, Squeegee, and Metacontam. Bars show the precision, recall, and F1 scores evaluated against ground-truth contaminant sets for the standard kit (**d**; *n* = 23 species) and microbiome kit (**e**; *n* = 7 species).

We evaluated decontamination performance by assessing whether contaminant removal reduced the distances between technical replicates, defined as samples collected from the same individual, body site, and date but processed with different kits (**Fig. 3b**). Compared with before decontamination, only Metacontam significantly reduced the Jaccard distances between technical replicates, whereas Decontam and Squeegee showed no significant improvement (**Fig. 3c**).

### Superior species-level decontamination against kit-specific ground truths

For each extraction kit, we defined the ground-truth (GT) contaminant species using the corresponding negative controls, yielding 7 GT species for the microbiome kit and 23 for the standard kit (**Supplementary Table 5**). Because the skin is continuously exposed to the environment and skin-associated taxa are commonly detected in indoor air and on surfaces, we conservatively identified GT species based on high-prevalence signals in negative controls^30,31^ (see **Methods**). We then benchmarked Metacontam against Decontam and Squeegee at the species level.

For the standard kit, Metacontam achieved precision/recall/F1 = 0.667 (10/15 species), 0.435 (10/23 species), and 0.526, outperforming both Decontam, which achieved 0.25 (1/4 species), 0.043 (1/23 species), and 0.074, and Squeegee, which achieved 0.25 (2/8 species), 0.087 (2/23 species), and 0.129 (**Fig. 3d**). For the microbiome kit, Metacontam achieved precision/recall/F1 = 0.357 (5/14 species), 0.714 (5/7 species), and 0.476, respectively, outperforming both Decontam, which achieved 0.235 (4/17 species), 0.571 (4/7 species), and 0.333, and Squeegee, which achieved 0.143 (2/14 species), 0.286 (2/7 species), and 0.190 (**Fig. 3e**). Overall, Metacontam consistently achieved the highest precision, recall, and F1 among the three methods, with F1 improvements of up to 7.1-fold for the standard kit and 2.5-fold for the microbiome kit, indicating the superior detection of kit-specific contaminants.

### Metacontam outperforms existing methods in low-biomass infant nasal metagenomes

We further validated Metacontam in a low-biomass setting by reanalyzing nasal shotgun metagenomes from healthy 24-month-old infants (n = 229), as reported by Zelasko et al^32^. GT species were defined as taxa with high prevalence across the 10 negative control samples, yielding 29 species (see **Methods** and **Supplementary Table 5**). At the species level, Decontam achieved precision/recall/F1 = 0.017 (24/1426 species), 0.828 (24/29 species), and 0.033; Squeegee achieved 0.111 (2/18 species), 0.069 (2/29 species), and 0.085; and Metacontam achieved 0.471 (16/34 species), 0.552 (16/29 species), and 0.508. Metacontam thus achieved an F1 score more than fivefold higher than that of Squeegee (Fig. 4a). To evaluate Squeegee more stringently, we additionally included oral samples from the same study to satisfy the requirement for distinct sample types; however, its performance further decreased, with precision/recall/F1 = 0.058 (3/52 species), 0.103 (3/29 species), and 0.074 (**Supplementary Fig. 5**).

**Fig. 4.**
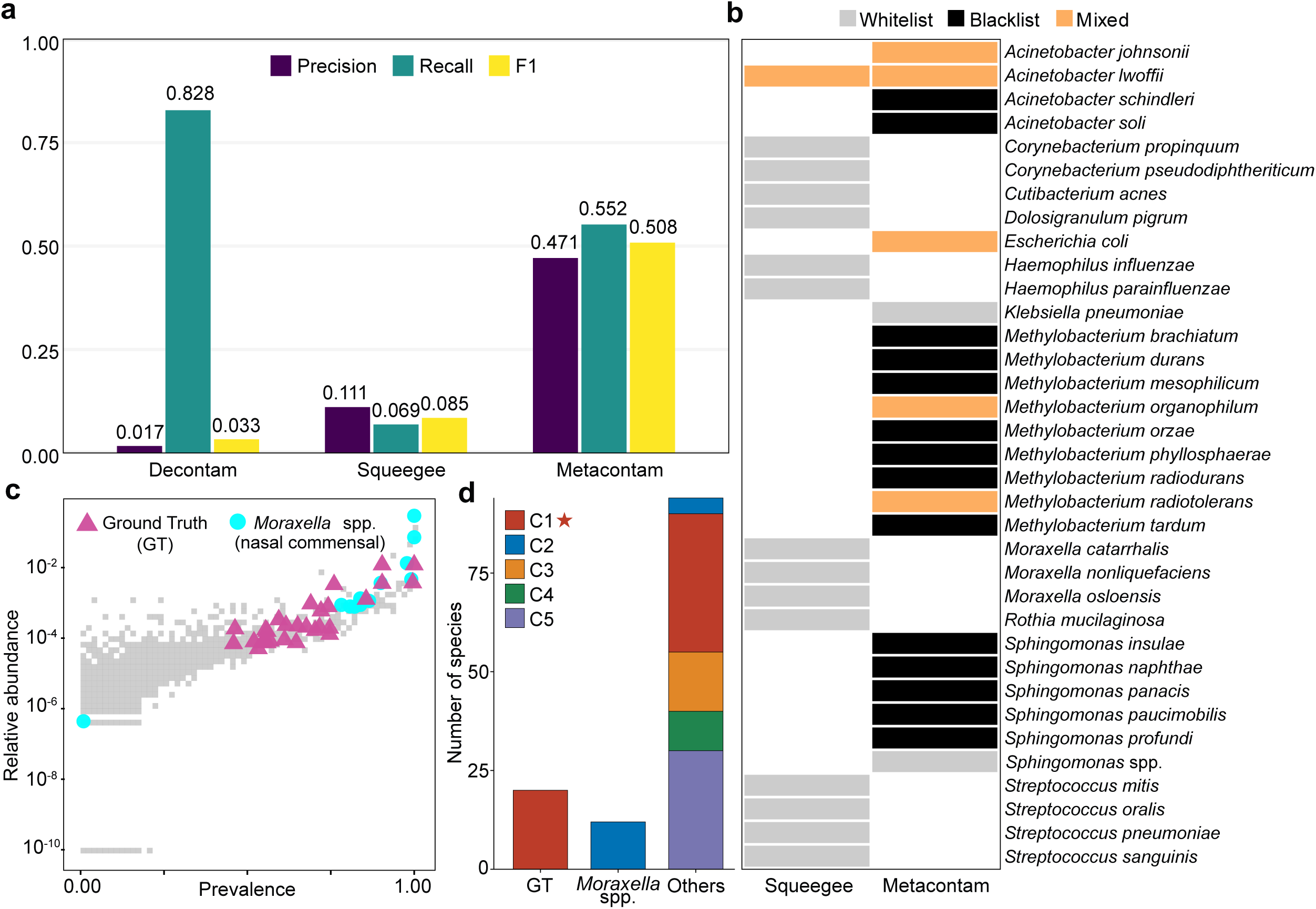
Performance on a nasal cohort and ecological specificity of predicted contaminants. **a**, Nasal shotgun metagenomes from healthy 24-month-old infants (*n* = 229). Species-level performance of Decontam, Squeegee, and Metacontam against the ground-truth contaminant set (29 species; see Methods). Bars show precision, recall, and F1 scores. **b**, Ecological specificity of predicted contaminants. Tile colors indicate Whitelist, Blacklist, or Mixed classification; species in neither list are omitted. **c**, Relationship between prevalence and mean relative abundance across samples. Moraxella spp. are shown as cyan circles, GT contaminants as purple triangles, and all other taxa are shown in grey. **d**, Number of species assigned to communities with ≥10 nodes, grouped into ground-truth (GT) contaminants, *Moraxella* spp., and other taxa. C1, the candidate contaminant community, is marked with a red star.

### Ecological specificity in infant nasal metagenomes

We next evaluated the ecological specificity of predictions by categorizing species according to their inclusion in a healthy nasal-associated whitelist, a contaminant-associated blacklist, or a mixed group containing species present in both lists (see Methods, “Whitelist and blacklist selection”; **Fig. 4b**). Metacontam predicted only two whitelist species as contaminants (2/34, 5.9%), in stark contrast to Squeegee, which incorrectly classified a large fraction of whitelist species as contaminants (14/18, 77.8%). Among the predicted species, Metacontam flagged 14 blacklisted species (14/34, 41.2%), whereas Squeegee flagged none (0/18, 0%). These results indicate that Metacontam not only improved the overall performance but also adopted an ecology-aware, conservative strategy by preferentially retaining whitelist taxa and more effectively flagging blacklist taxa.

*Moraxella* spp. is a representative example of this ecological distinction. In the prevalence-abundance space, GT species largely occupied the high-prevalence, high-abundance region, but *Moraxella* spp. also fell within this range, which is consistent with their status as common upper airway residents^33,34^ (**Fig. 4c**). This indicates that even highly prevalent and abundant taxa are not necessarily contaminants. Consistent with this observation, Squeegee, which relies primarily on the prevalence and coverage, misclassified *Moraxella* spp. as contaminants. In contrast, Metacontam correctly classified them as non-contaminants and placed GT and *Moraxella* spp. in distinct network communities (C1 and C2, respectively; **Fig. 4d**).

### Metacontam improves classification accuracy in challenging low-biomass tissue microbiomes

Although tissue microbiome samples have inherently low biomass and are susceptible to contamination, patient-matched comparisons of tumors and adjacent normal tissues have revealed distinct microbial compositions in multiple cancer types^6,18,35,36^. Motivated by these observations, we investigated whether decontamination improved the classification performance. We compared four preprocessing strategies: no decontamination, Decontam, Squeegee, and Metacontam. As shown in **Fig. 5a**, the models were trained on TCGA colorectal tissue whole-genome sequencing data (*n* = 173) and evaluated on an independent non-TCGA external cohort^37^ (*n* = 28), with contaminant removal performed separately for each center. In the external cohort, Metacontam achieved the highest AUC (0.770), outperforming no-decontamination (0.638), Decontam (0.638), and Squeegee (0.709) (**Fig. 5b**); it also ranked highest in the training set (**Supplementary Fig. 6a**).

**Fig. 5.**
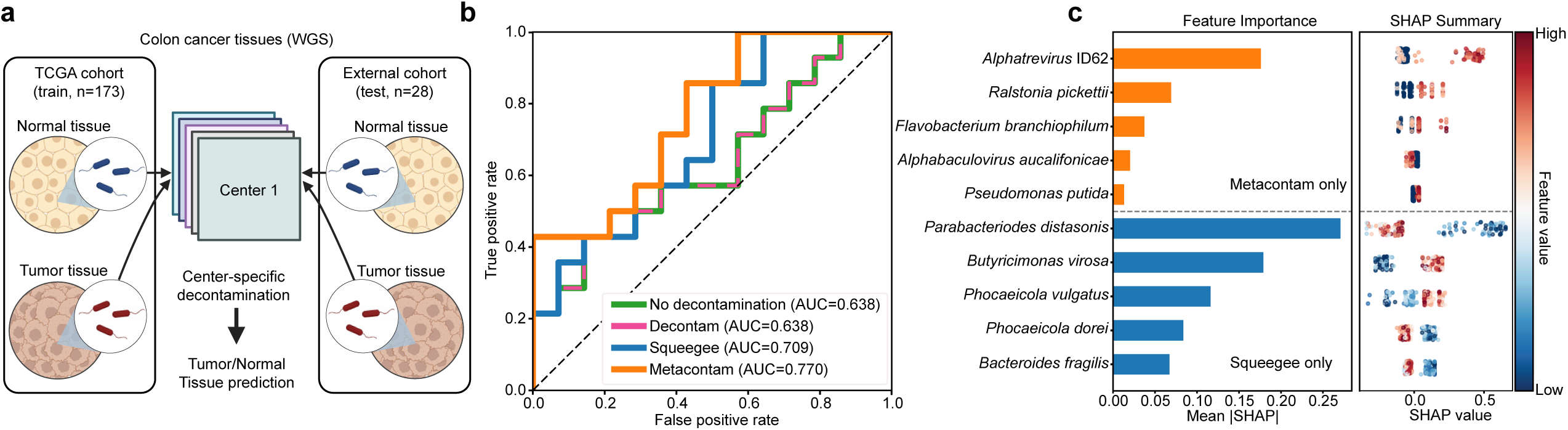
Metacontam improves discrimination between tumor and normal tissues in colorectal cancer (CRC). **a**, Overview of the CRC analysis workflow using microbiome reads from whole-genome sequencing data of colorectal tissue. The models were trained on the TCGA cohort (*n* = 173; normal = 24, tumor = 149) using no decontamination, Decontam, Squeegee, or Metacontam preprocessing, and evaluated on an independent non-TCGA cohort (*n* = 28; normal = 14, tumor = 14). **b**, ROC curves of the independent external (non-TCGA) test cohort. The no-decontamination and Decontam curves overlap. **c**, SHAP-based interpretation of model features. Taxa predicted as contaminants only by Metacontam or only by Squeegee are compared. Left: mean |SHAP| computed from the original feature matrix for the top six Metacontam-only and Squeegee-only taxa. Right: SHAP beeswarm plots for the same taxa (positive values shift predictions toward tumor; negative values toward normal).

Next, we examined the taxa removed using each method and their feature importance. Metacontam flagged *Ralstonia pickettii* and *Flavobacterium branchiophilum* as contaminants, both of which belong to genera commonly detected in environmental sources, whereas Squeegee-flagged *Parabacteroides distasonis* and *Butyricimonas virosa* belong to genera that are typical gut commensals that have been linked to colorectal cancer^38–41^. Among the top six most important features selected under each preprocessing strategy, taxa uniquely retained by Metacontam generally showed lower feature importance than those unique to Squeegee, with *Alphatrevirus* being the only exception. This phage may reflect co-introduction with contaminant *E. coli* or technical artifacts, such as PhiX-related contamination, as reported in the TCGA sequencing data (**Fig. 5c**)^42,43^. We further evaluated two additional tissues without external cohorts. In the esophagus, Metacontam performed slightly better than Decontam and the no-decontamination baseline, whereas in the stomach, the methods performed similarly (**Supplementary Fig. 6c,d**). Overall, these results suggest that Metacontam effectively mitigates contamination-related signals while preserving biologically informative signals.

### Robustness of Metacontam across varying sample sizes and contamination levels

Using a real fecal metagenome dataset from Pomyen et al.^44^, we simulated contamination to assess the stability of Metacontam across sample sizes and contamination levels. At a fixed contamination level, the performance was similar at *n* = 50 and *n* = 30 (F1 = 0.893 and 0.889, respectively) but dropped at *n* = 10 (F1 = 0.568) (**Supplementary Fig. 10a**). Next, we varied the contamination levels using a fixed sample size (*n* = 50). F1 remained high at 1% and 0.5% spike-in (0.873 and 0.893, respectively) but decreased at 0.1% spike-in (0.625) (**Supplementary Fig. 10b**). At a 0.1% spike-in, the average read count per contaminant species was below 33, and several ground-truth species exhibited abrupt ANI drops (**Supplementary Fig. 2**). Overall, Metacontam was stable across moderate sample sizes and contamination levels, with its performance degrading primarily under extremely low-signal conditions.

### Experimental validation of Metacontam using fecal-DNA spike-in into low-biomass mouse BALF samples

We further evaluated our core assumptions using a controlled mouse experiment designed to mimic the contamination of a low-biomass respiratory sample. Bronchoalveolar lavage fluid (BALF) was collected from eight individual mice, and contamination was simulated by spiking fecal DNA from a single mouse into all BALF DNA samples (**Supplementary Fig. 9a**). BALF- and fecal-enriched species formed well-separated groups in the correlation network, with known respiratory tract commensals such as *Pasteurella multocida* appearing among the BALF-enriched taxa (**Supplementary Fig. 9b–c**). Moreover, fecal-enriched species showed significantly higher ANI values than BALF-enriched species (**Supplementary Fig. 9d**). Although Metacontam was not directly applied to this dataset due to the lack of a mouse-specific blacklist, these results provided experimental support for our core hypothesis that contaminants and genuine taxa are separable by network structure and genomic similarity (ANI).

## Discussion

Low-biomass shotgun metagenomic samples are particularly vulnerable to reagent and laboratory contamination, and even trace amounts of exogenous DNA can overwhelm the true signal. Yet in many datasets, negative controls are unavailable, limiting widely used approaches such as Decontam. Squeegee operates without negative controls but requires compositionally distinct sample types and has not been validated on datasets composed exclusively of low-biomass samples.

To address these limitations, we present Metacontam, a control-free decontamination method that leverages network structure and genome-wide similarity. Compared to Decontam and Squeegee, Metacontam achieved the highest F1 score across diverse low-biomass metagenomic datasets, recovering low-abundance and low-prevalence contaminants while retaining ecologically relevant taxa and remaining robust across varying sample sizes and contamination levels.

Metacontam specializes in low-biomass datasets. In low-biomass samples, contaminants constitute a significant share of the reads, enabling reliable abundance estimates and ANI-based refinement. Consistent with this, Metacontam significantly decreased Shannon and Simpson diversity in placental and breast milk samples, indicating its clearest downstream effect in low-biomass profiles (**Supplementary Fig. 3a–b**). In contrast, high-biomass samples are dominated by true taxa, which obscured contaminant signals and yielded sparse contaminant coverage, degrading both abundance and ANI accuracy. Consistent with this, in an evaluation using only oral metagenome data (a high-biomass context), Metacontam achieved an F1 of 0.20, compared with 0.12 for Squeegee and 0.01 for Decontam (**Supplementary Fig. 5**). Although Metacontam performed the best, all the methods performed poorly on datasets composed solely of high-biomass samples.

We constructed a blacklist based on the taxa frequently reported as contaminants in human metagenomic studies. Building on this baseline, one way to enhance Metacontam’s performance is to adapt the blacklist to the study context: add species that are ecologically implausible for the environment and those known to recur as laboratory or sequencing-center contaminants, and remove species that are plausible for the environment.

Our validation focused on shotgun metagenomic datasets because the refinement step relies on ANI, which is not well-suited to 16S rRNA amplicon data. Nevertheless, across all validation datasets, using only the blacklist and seeded Louvain algorithm produced F1 scores of at least 0.36. Therefore, for 16S rRNA datasets, omitting ANI is a practical option, although its performance may be lower than that of the full Metacontam pipeline, which includes the ANI refinement step.

Finally, a major advantage of Metacontam is that the key parameters are automatically set from the data. In Squeegee, parameters such as prevalence and minimum read count are fixed at default values, and there is no guidance for adapting them to a dataset. Even for Decontam, the optimal parameters varied across studies^4,13^. In contrast, in Metacontam, both the prevalence threshold and the ANI threshold used for the final contaminant calls are tuned to the dataset. This automation reduces the user burden and bias and is expected to provide optimal performance for each dataset.

In summary, Metacontam provides a robust, control-free framework for decontaminating shotgun metagenomic data, particularly in low-biomass settings, by integrating network-based community detection with ANI-based refinement, offering improved performance over existing methods without requiring negative controls.

## Methods

### Whitelist and blacklist selection

We built body site-specific healthy whitelists from the mBodyMap as follows: Skin from the Healthy-Skin section, Oral from Healthy-Oral, and Nasal from the union of Healthy-Nose and Healthy-Upper respiratory tract. We retained mBodyMap’s default inclusion criteria (species observed in ≥ 2 healthy samples and with median relative abundance ≥ 0.01%)^45^.

Following Salter et al., we curated a set of blacklist species commonly found in negative controls: 1,781 species across 31 genera, each reported in at least two independent studies^46^. Each whitelist was filtered against a blacklist before downstream use. The final whitelist sizes after blacklist filtering were skin 811/874 (removed 63), nasal 958/1034 (removed 76), and oral 350/351 (removed 1) (kept/total; removals in parentheses).

### Ground truth construction

Ground-truth contaminants were defined as taxa whose prevalence in negative controls exceeded a dataset-specific threshold. Thresholds were derived exclusively from the negative controls for each dataset (skin, standard kit *n* = 23; skin, microbiome kit *n* = 36; nasal *n* = 12; oral *n* = 11). A species was considered present in a sample if its relative abundance was ≥ 0.001, and prevalence was the fraction of negative-control samples in that dataset where it was present.

For each dataset, we scanned prevalence thresholds from 0.10 to 1.00 in increments of 0.10; at each threshold, we computed the whitelist ratio, which is the proportion of retained species that also appeared in the corresponding healthy body-site whitelist. The prevalence threshold was defined as the cutoff with the lowest whitelist ratio; the selected thresholds are indicated by the vertical dashed lines in **Supplementary Fig. 7a–d**. Using this procedure, the selected prevalence thresholds and contaminant counts were: Skin: microbiome kit 0.60 (7 species), Skin: standard kit 0.40 (23 species), Nasal 0.80 (29 species), and Oral 1.00 (14 species). For the maternal-infant dataset, we did not re-estimate thresholds; instead, we adopted the permissive ground-truth set from the Squeegee study (Liu et al., 2022)^16^, comprising 31 species from 16 genera, curated from multiple negative controls with manual removal of known false positives.

### Preprocessing and abundance profiling

All raw metagenomic sequencing reads were quality controlled using fastp (v0.23.4; options: --dedup --low_complexity_filter --detect_adapter_for_pe) for adapter trimming and removal of duplicate or low-complexity reads^47^. Host-derived sequences were removed by mapping against the CHM13v2.0 human reference genome using Bowtie2 (v2.3.5.1, default parameters), and only read pairs in which both mates were unmapped were retained using samtools (v0.1.19)^48–50^.

For the tissue microbiome datasets, we performed more stringent human read removal to account for the high host DNA fraction. Two pre-processing pipelines were applied.

i. fastp, followed by host removal against CHM13v2.0, and
ii. Trimmomatic (v0.39; ILLUMINACLIP:2:30:7 MINLEN:50 TRAILING:20 AVGQUAL:20 SLIDINGWINDOW:20:20) and BBDuk (entropy filter, threshold ≥ 0.6; part of the BBMap suite, Bushnell B., sourceforge.net/projects/bbmap/) followed by host removal against GRCh38^48,51^.

Taxonomic profiling was performed using Kraken2 (v2.1.3) in paired-end mode using the standard database built in August 2023 (kraken2-build --standard). This database contains complete bacterial, archaeal, and viral genomes from RefSeq, human reference genome, and UniVec_Core sequences. Species-level abundance estimates were refined using Bracken (v2.9) at the species rank^52,53^.

### Network construction and community detection

Microbial association networks were constructed using NetCoMi (v1.1.0)^54^. After multiplicative-replacement zero imputation and CLR transformation, we estimated pairwise Pearson correlations between taxa and retained only positive links with r ≥ 0.45. The community structure was subsequently inferred using a Louvain-based procedure that extended the standard algorithm by introducing a seeded initialization step in which all blacklist species were grouped into a single community before optimization. During this seeding step, edges among blacklist nodes followed the standard Louvain edge-handling rules: intra-community edges were converted to self-loops, inter-community edges were aggregated as weighted edges in the coarse-grained graph, and standard modularity optimization was then applied^28^.

### Adaptive prevalence-threshold estimation

For each taxon *t*, prevalence *p_t_* was defined as the fraction of samples with read counts ≥ *r*_min_(default *r*_min_ = 2). From the blacklist taxa detected in the dataset, we selected the *N* = 100 most prevalent taxa and defined *p*_black_ as the median prevalence. To quantify overall association strength, we constructed a Pearson correlation network using the *N* = 100 most prevalent taxa (irrespective of blacklist membership) and defined *c*_med_ as the median positive pairwise correlation coefficient (edges with *r* ≥ 2). Let n be the number of samples.

Shift term:

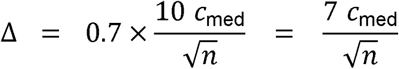

Prevalence threshold:

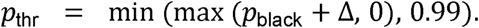

Post-hoc adjustment: If applying *p*_thr_ retained ≥ 500 taxa, we incremented *r*_min_ by 1 and recomputed *p_t_*, *p*_black_, *c*_med_ and *p*_thr_ until fewer than 500 taxa remained. Threshold estimation requires at least one blacklist taxon to be present in the data.

### ANI-Based Identification of Contaminant Species

The consensus ANI (conANI) values were calculated using InStrain (v1.9.1) for species with complete reference genomes. For each species, the mean conANI was computed across sample pairs, considering only the alignment regions with >50 bp of the compared bases^55^.

To reduce the computational burden of exhaustive pairwise comparisons, Mash (v2.1) was applied for stratified sampling. Mash distances were computed from the read sketches (k=21, sketches =100,000). If the number of possible sample pairs exceeded 1,000, 1,000 pairs were selected across 10 Mash distance bins; otherwise, all sample pairs were used^56^.

### ANI threshold for final prediction

Let µ_s_ denote the species-level mean conANI.

With S as the set of species having µ_s_ and B as the blacklist, define the blacklist ratio.

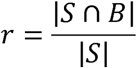

Using base percentile *p*_0_ = 0.60, set the effective percentile

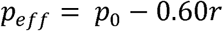

and the ANI threshold

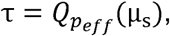

Where *Q_P_* is the *p*-th percentile of {µ_s_ :*s* ɛ *S*},

A species is classified as a contaminant if µ_s_ ≥ τ; otherwise it is classified as a non-contaminant.

**Design note**. A higher proportion of blacklisted species lowers the ANI threshold, thereby yielding more contaminant calls.

### Tissue microbiome decontamination and classification

For decontamination, we used a center-specific strategy; taxa predicted to be contaminants by Metacontam or Squeegee were removed only from the samples sequenced at that center. Cohorts covered multiple centers and tissues. Colorectal cancer: Baylor College of Medicine (BCM, n=107), Harvard Medical School (HMS, n=66), and University of Birmingham (UoB, n=14); stomach: HMS (n=141), MD Anderson (MDA, n=5); esophagus: MDA (n=43), Washington University School of Medicine (WUSM, n=20). BCM and HMS comprised multiple tissue types, satisfying Squeegee’s requirements for distinct sample sets.

XGBoost was used for classification^57^. The cancer models were trained using TCGA and evaluated once in an independent non-TCGA test cohort. In contrast, the esophageal and stomach models used TCGA only, with performance estimated by 5-fold cross-validation. We performed a grid search over the feature set size (top taxa by mean abundance: 100-3000 in steps of 100) and XGBoost hyperparameters (max_depth ∈ {3, 5}, n_estimators ∈ {100, 300}, learning_rate = 0.1, subsample = 1). The combination with the highest 5-fold cross-validated AUC in TCGA was selected.

For colorectal cancer, the selected configuration was retrained on the full TCGA training set and then evaluated in an external cohort.

### Metagenomic simulation data generation

Thirty contaminant species were randomly chosen from the curated blacklist, with one species per genus. For each species, one RefSeq complete genome (excluding plasmids) was retained, and contaminant reads were generated with InSilicoSeq (v1.5.4) using the HiSeq model^58^. For each simulated sample, a log-normal composition across the 30 species was used to define a common base ratio among contaminant species, and slight sample-specific variation was introduced by sampling from a Dirichlet distribution (α = base x 1e5), keeping relative ratios nearly preserved across samples. Native fecal reads were retrieved from real datasets, read pairs containing any ‘N’ were removed, and reads were randomly subsampled. Simulated contaminant reads were spiked into each sample at proportions of 0.1%, 0.5%, or 1% of the total reads to produce the final simulation datasets.

### Mouse lung microbiome DNA preparation

Eight- to 10-week-old C57BL/6J mice were used for the experiments described in this study. The experiments were conducted according to protocols approved by the Institutional Animal Care and Use Committee of the Gwangju Institute of Science and Technology (GIST). Mice were bred and maintained under specific pathogen-free conditions at GIST. To collect bronchoalveolar lavage fluid (BALF), 1 ml of ice-cold Ca2+-free PBS was injected into the trachea using a 22G catheter. After instillation and withdrawal two times, BALF cells were collected by centrifugation at 15,000 x g for 15 min at 4c. DNA from the lung microbiome was extracted from cell pellets using the LaboPassTM Tissue Genomic DNA Isolation Kit Mini (CosmoGeneTech, Korea) according to the manufacturer’s instructions.

### Decontamination with Decontam and Squeegee

For the skin, nasal, and oral benchmark datasets, Decontam (v1.24.0) was applied in prevalence mode with matched negative controls. Following the original Squeegee benchmarking procedure, species with at least 30 reads or a relative abundance of at least 0.0005 were retained to construct the abundance table, and contaminants were identified using a threshold of 0.01. For tissue microbiome datasets, Decontam (v1.24.0) was applied in frequency mode with default parameters using the DNA concentration values for each sequencing center. Squeegee (v0.1.3) was run on the same taxonomic profiles using the default scoring parameters.

## Data availability

Sequencing reads generated in the mouse BALF fecal DNA spike-in experiment are available in the NCBI SRA under BioProject accession PRJNA1424585. Benchmarking datasets are publicly available under BioProject accession numbers PRJNA725597 (maternal infant), ERP124610 (skin extraction kit comparison dataset; Qiita study 12201), PRJNA1078287 (infant nasal, oral), and PRJNA932948 (fecal dataset used for simulation). The TCGA WGS data were accessed via the NCI Genomic Data Commons under controlled access through dbGaP (study accession number phs000178). Sample-level metadata are provided in **Supplementary Table 3**.

## Code availability

The Metacontam implementation is available at https://github.com/iamjunwoojo/Metacontam. The version used in this study corresponds to release version 0.0.1. The code for benchmarking, analysis, and figure generation is available at https://github.com/iamjunwoojo/Submission_figure_Metacontam with reproducible environments provided in the repository.

## Contribution to Authorship

Sun.L. and J.C. supervised the study. J.J. and Sun.L. wrote the manuscript. J.J. conceived the study and performed the bioinformatics analyses. H.L. conducted the mouse experiments. J.W.B. contributed to the early development of the study concept. Su.L. contributed to data interpretation and visualization. Sun.L., J.C., S.S., A.M., V.S. reviewed the manuscript and provided critical comments. All authors approved the final version of the manuscript.

