## Supplementary material for "Metacontam: A Negative Control-Free Decontamination Method for Metagenomic Analysis": Supplemtary Figures

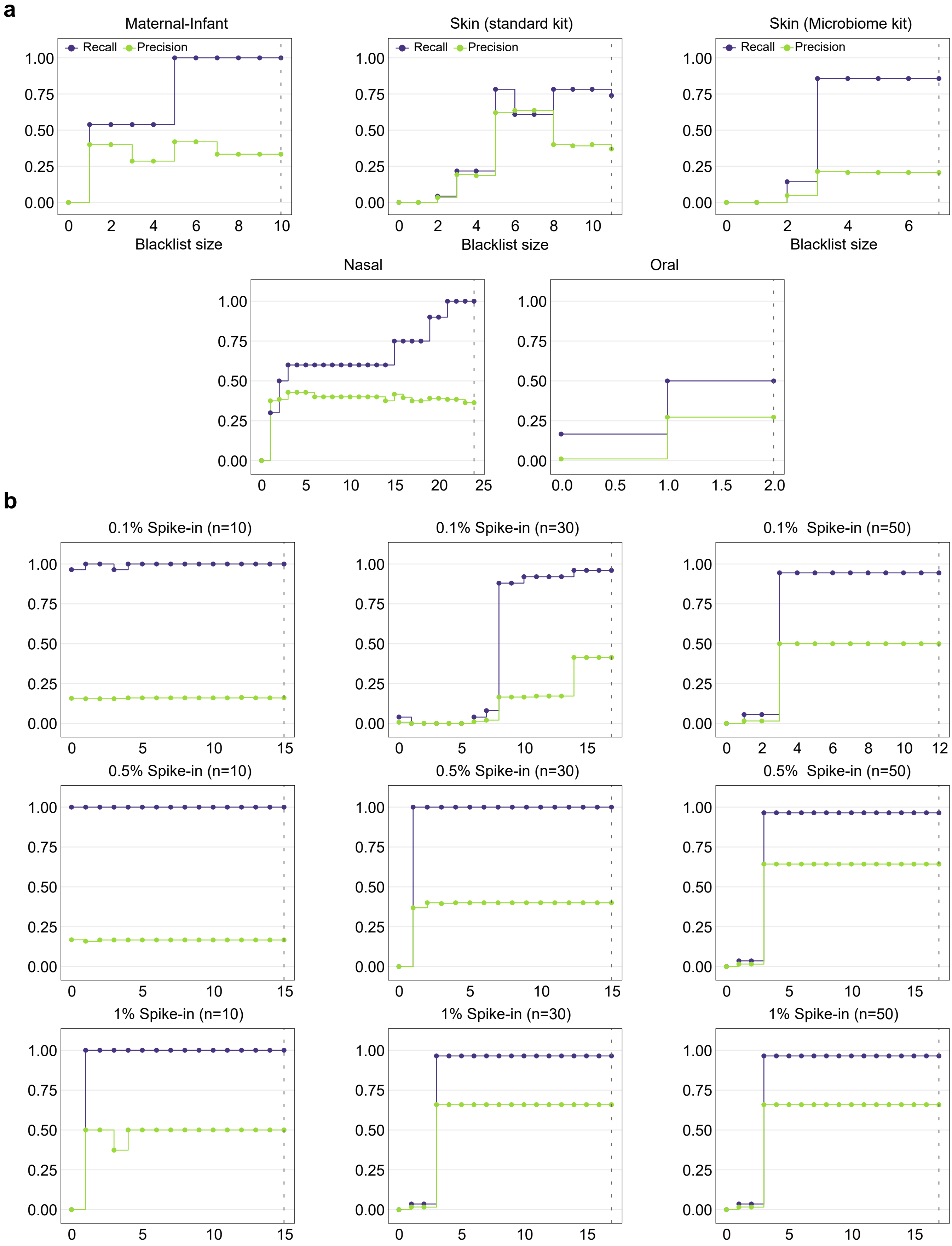


**Supplementary Fig. 1 | Recall and Precision of seeded Louvain community detection across blacklist sizes.**
Each panel shows Recall and Precision of the candidate contaminant community as blacklist size increases from 0 to N. **a,** Upper panels correspond to five real datasets (MI, Standard, Microbiome, Nasal, Oral); **b,** lower panels correspond to simulated datasets across three contamination levels (0.1%, 0.5%, 1%) and three sample sizes (n = 10, 30, 50). Recall is defined as the proportion of ground-truth (GT) species recovered within the contaminant community; Precision is defined as the proportion of contaminant community members that are GT species. The dotted vertical line indicates the full blacklist size (N) for each dataset.

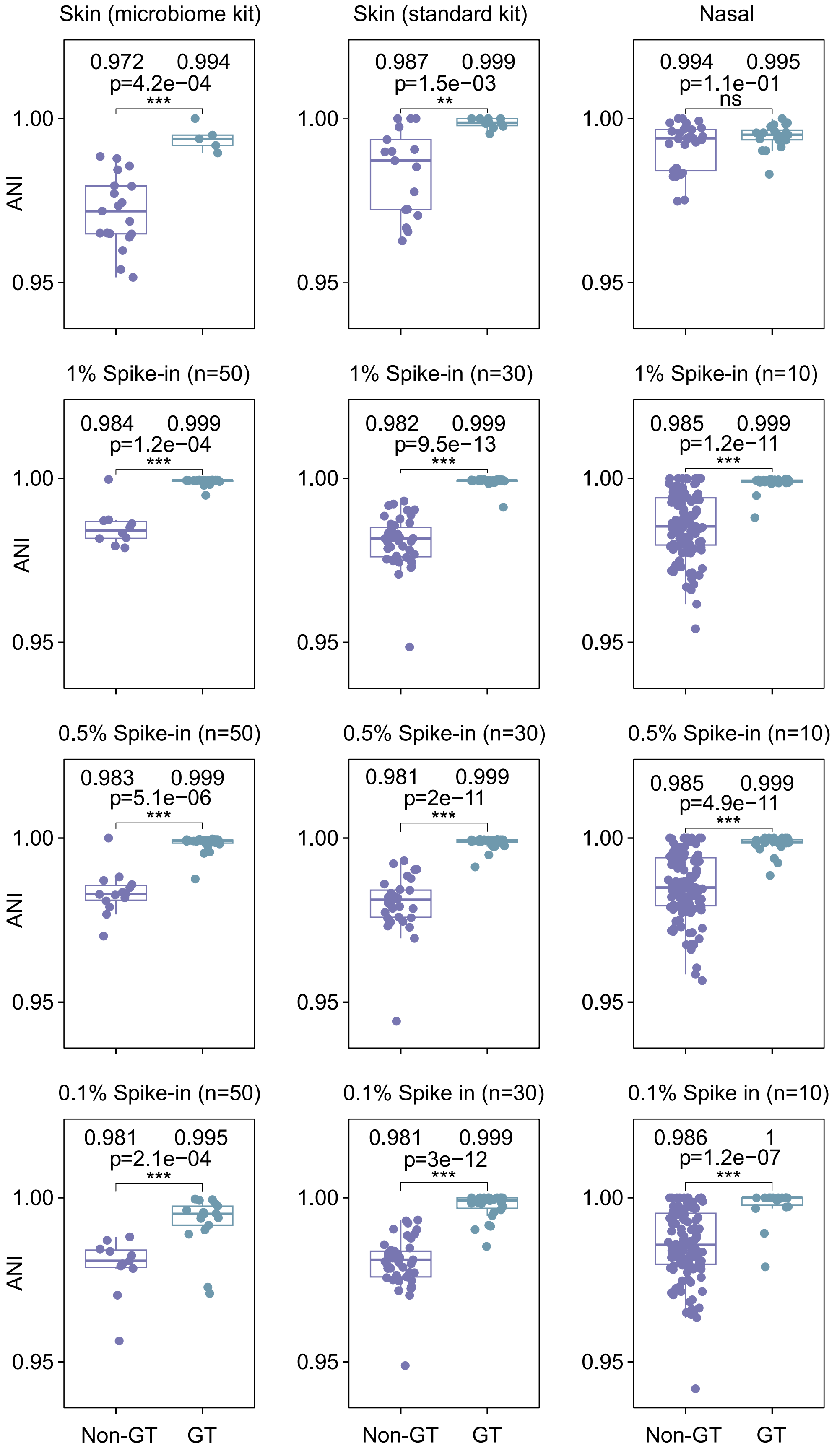


**Supplementary Fig. 2 | ANI within candidate contaminant communities.**
Each panel shows the per-species mean ANI for taxa in the candidate contaminant community for each dataset and condition. Species are labeled as GT or non-GT. Values above boxes indicate medians. One-sided Wilcoxon rank-sum tests evaluated whether GT > non-GT; * $p \leq 0.05$; ** $p \leq 0.01$; *** $p \leq 0.001$; n.s., $p > 0.05$.

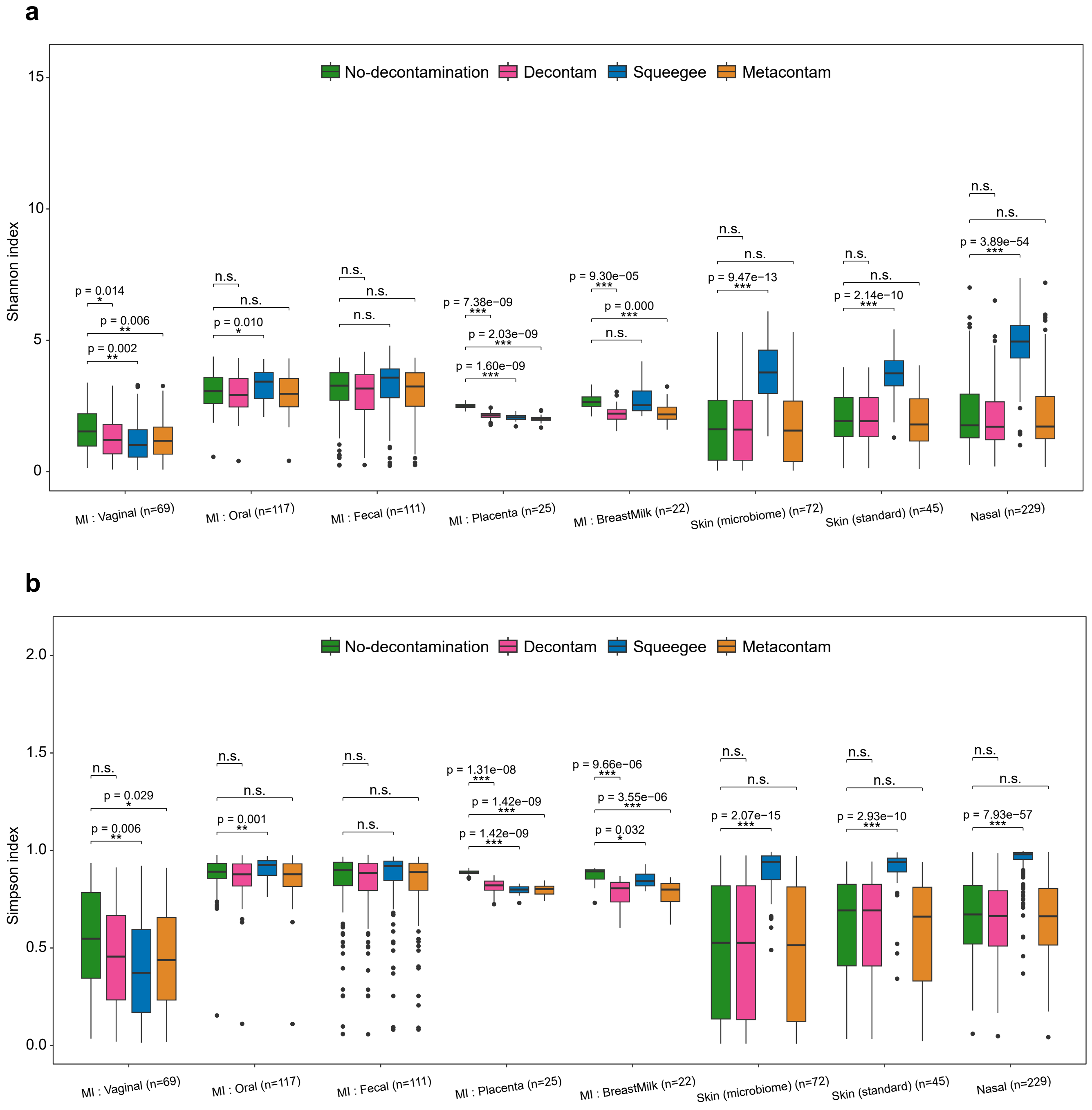


**Supplementary Fig. 3 | Alpha-diversity after removal of predicted contaminants across datasets.** **a,b,** Shannon's index **(a)** and Simpson's index **(b)** were calculated from species-level relative abundances for Maternal-Infant datasets, Skin (standard kit), Skin (microbiome kit) and Nasal groups. Samples were evaluated under No-decontamination, Squeegee (Squeegee-predicted taxa removed) and Metacontam (Metacontam-predicted taxa removed). Boxplots summarize distributions (center line, median; box, IQR; whiskers, 1.5×IQR). Two-sided Wilcoxon rank-sum tests within each group. * $p \le 0.05$; ** $p \le 0.01$; *** $p \le 0.001$; n.s., $p > 0.05$.

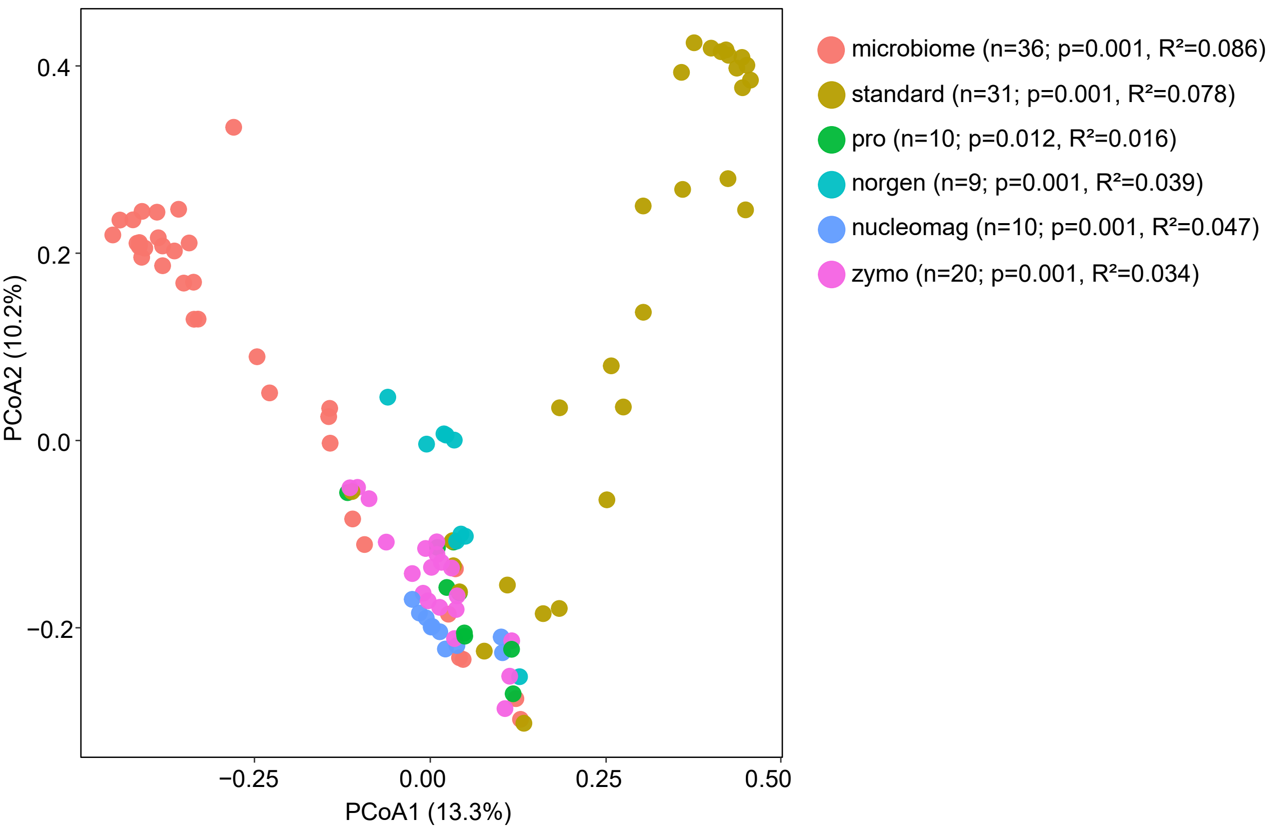


**Supplementary Fig. 4 | PCoA of negative controls across extraction kits in the skin validation dataset.**

Principal coordinates analysis (PCoA) of Bray-Curtis dissimilarities computed from species-level relative abundances for sterile water blank samples. Points represent samples and are colored by extraction kit (microbiome = MagMAX Microbiome Ultra; standard = Qiagen PowerSoil; pro = Qiagen PowerSoil Pro;norgen = Norgen Stool; nucleomag = Macherey-Nagel NucleoMag Food; zymo = ZymoBIOMICS 96 MagBead). The legend reports one-vs-other PERMANOVA results (vegan::adonis2, 999 permutations) for each kit, including sample count (n), p-value and $R^2$.

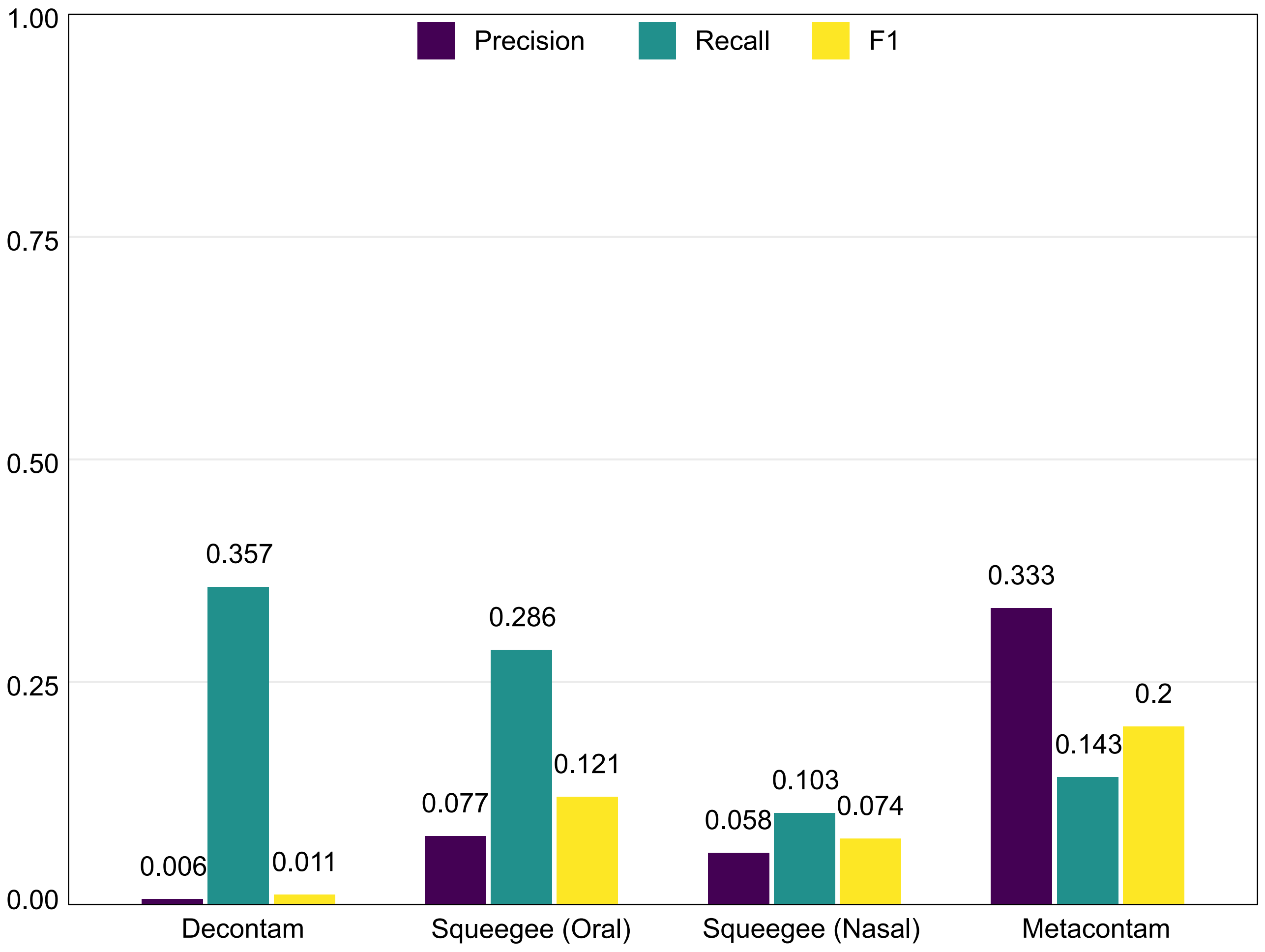


**Supplementary Fig. 5 | Species-level benchmarking using oral- and nasal-derived ground-truth contaminant sets.**

Decontam and Metacontam were applied to oral samples and evaluated against an oral negative-control-derived ground-truth contaminant set (14 species). Squeegee was run on pooled nasal and oral samples (n = 441; nasal = 229; oral = 212). Squeegee (Oral) was evaluated against the oral ground-truth set, and Squeegee (Nasal) against the nasal ground-truth set (29 species).

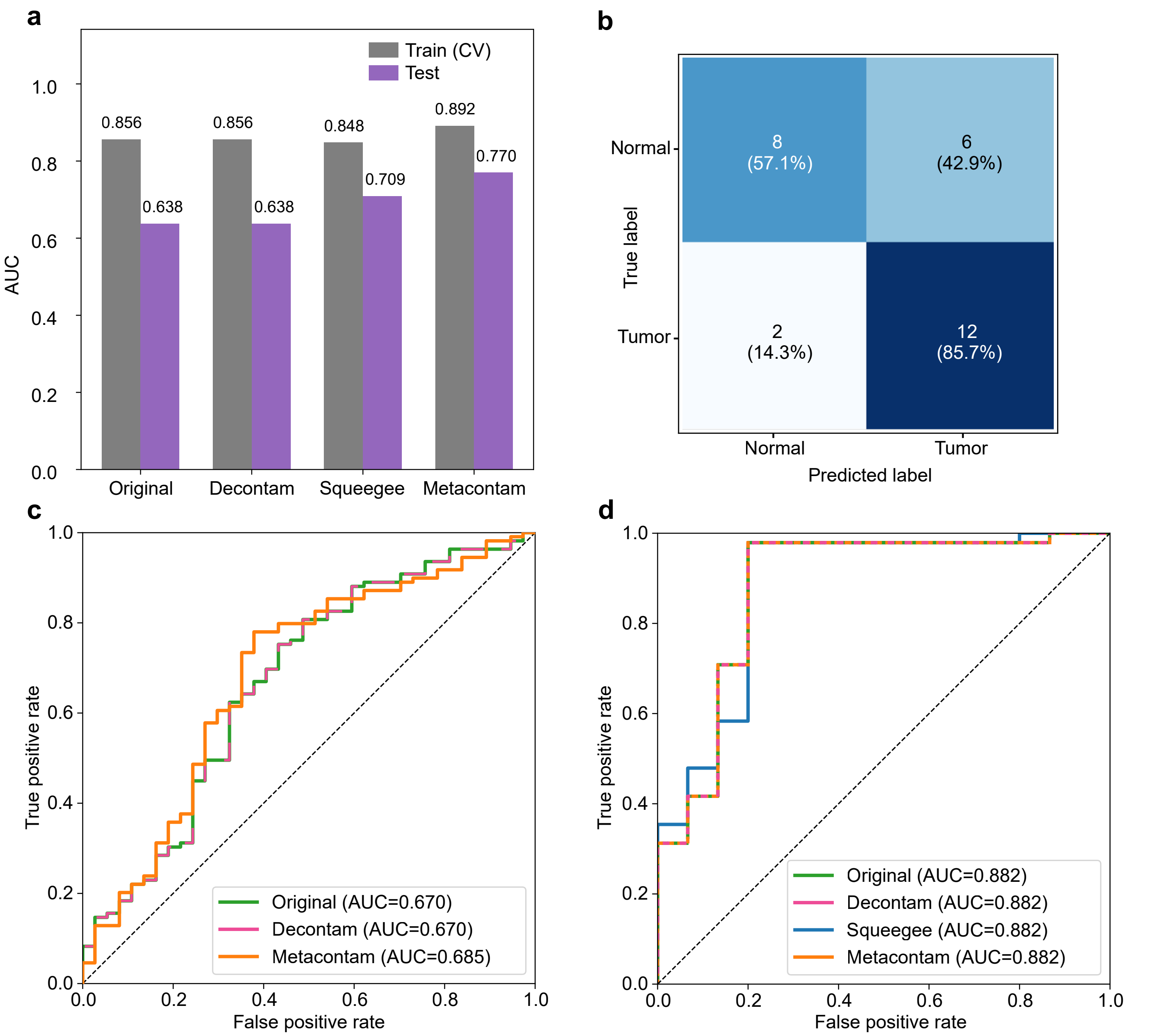


**Supplementary Fig. 6 | Tumor–normal tissue classification performance after decontamination.** **a,** Colon: AUC comparison across preprocessing methods; bars show cross-validated training AUC and external test AUC. XGBoost model used the top-N taxa selected by mean relative abundance in the training set. **b,** Colon: Confusion matrix for the Metacontam model. Cells show counts with row-normalized percentages. The decision threshold was selected to maximize PR-F1 on out-of-fold predictions and applied unchanged to the test set. **c,d,** Esophagus **(c)** and stomach **(d)**: ROC curves for the best-performing configuration per preprocessing method. AUCs were computed from out-of-fold predictions using 5-fold stratified cross-validation.

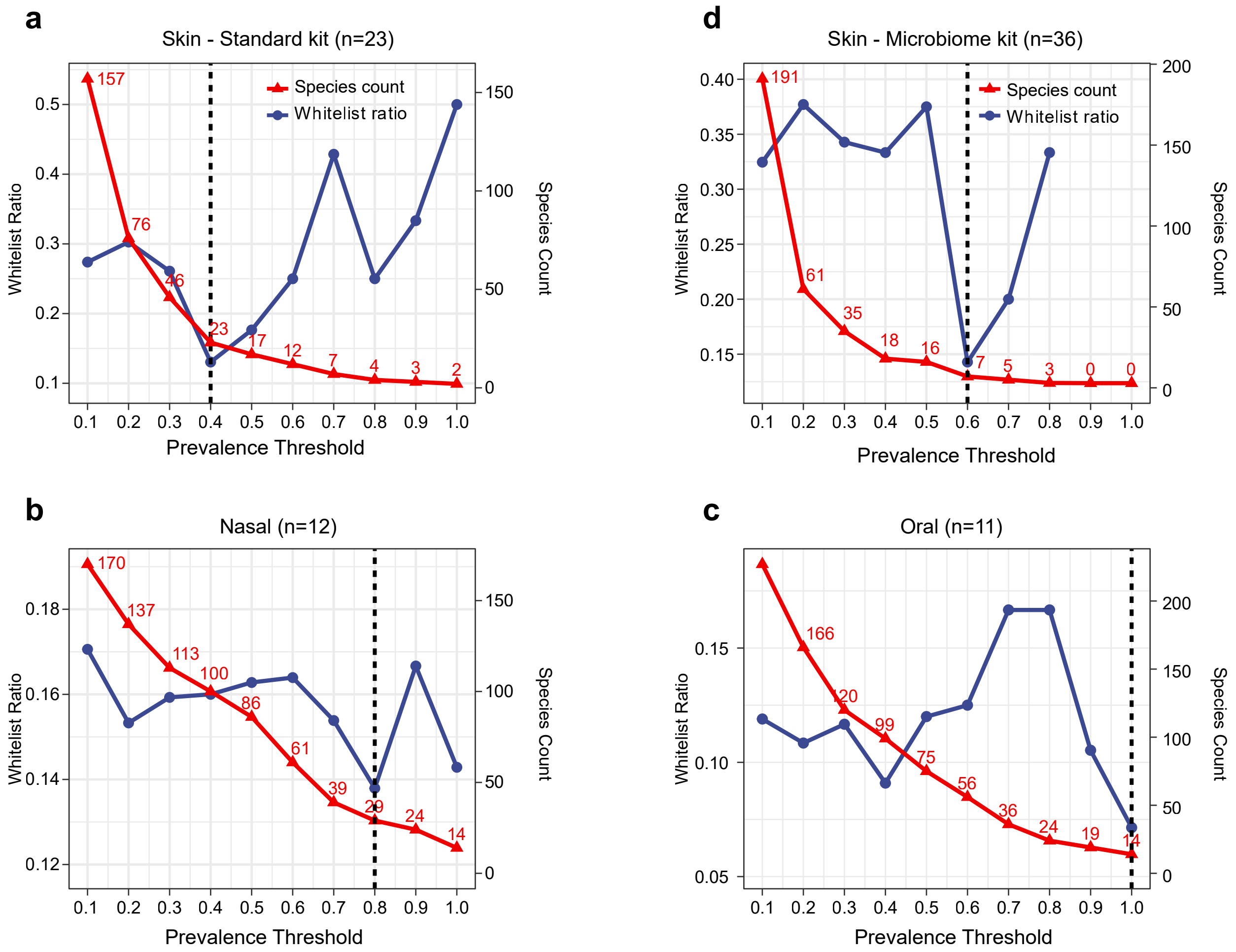


**Supplementary Fig. 7 | Ground-truth contaminant thresholds from negative controls.**
**a-d,** For prevalence thresholds from 0.1 to 1.0, we plotted the number of species passing each threshold (red line) and the whitelist ratio (blue line). Prevalence was the fraction of negative controls with species relative abundance ≥ 0.001. The dashed line marks the threshold with the lowest whitelist ratio, adopted as the groundtruth contaminant threshold. Negative-control sample sizes: **a,** Skin-Standard (n = 23); **b,** Skin-Microbiome (n = 36); **c,** Nasal (n = 12); **d,** Oral (n = 11).

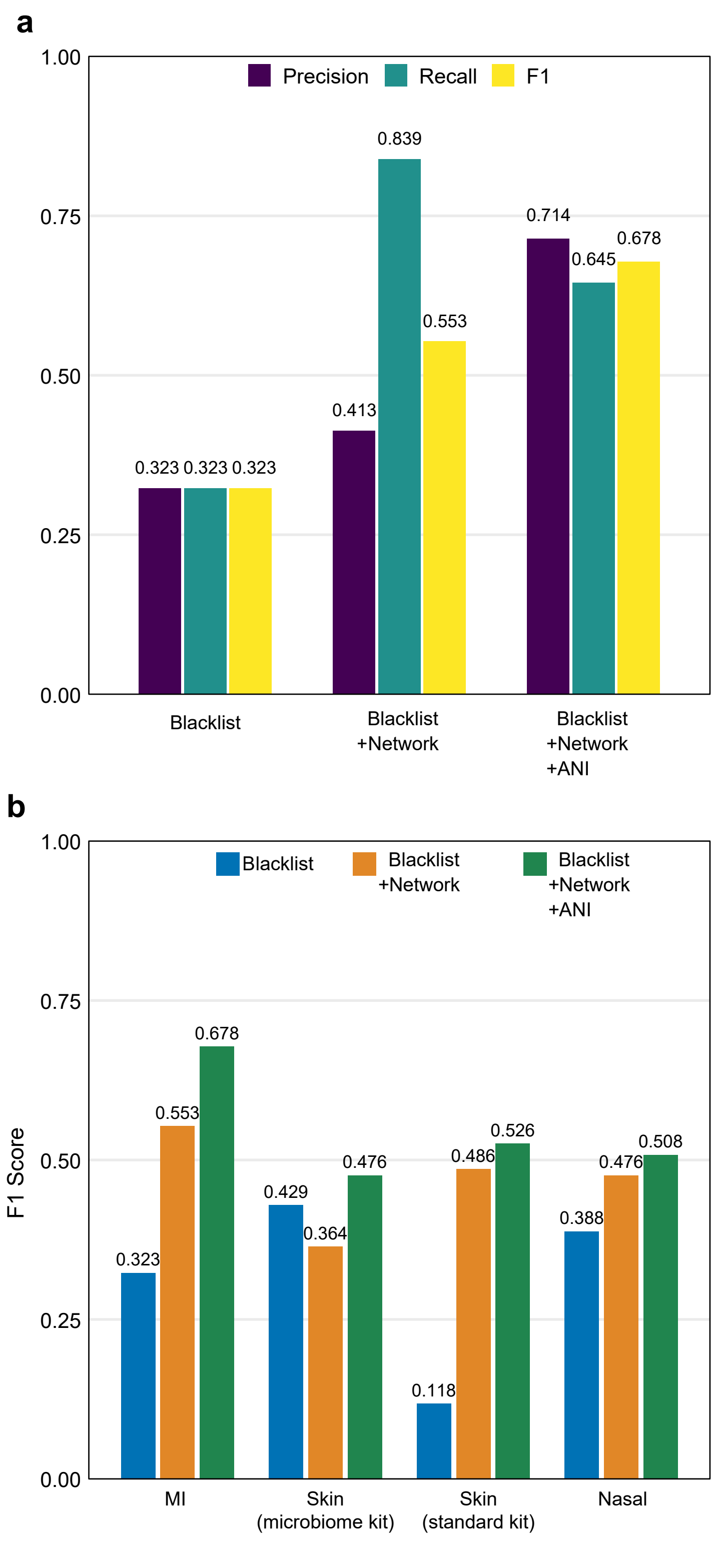


**Supplementary Fig. 8 | Stepwise evaluation across contamination signatures.**
F1 scores across datasets under three strategies: blacklist only, blacklist + network, and blacklist + network + ANI. **a,** Maternal–infant dataset. **b,** Four low-biomass sample types, including maternal-infant (MI), skin, and nasal samples.

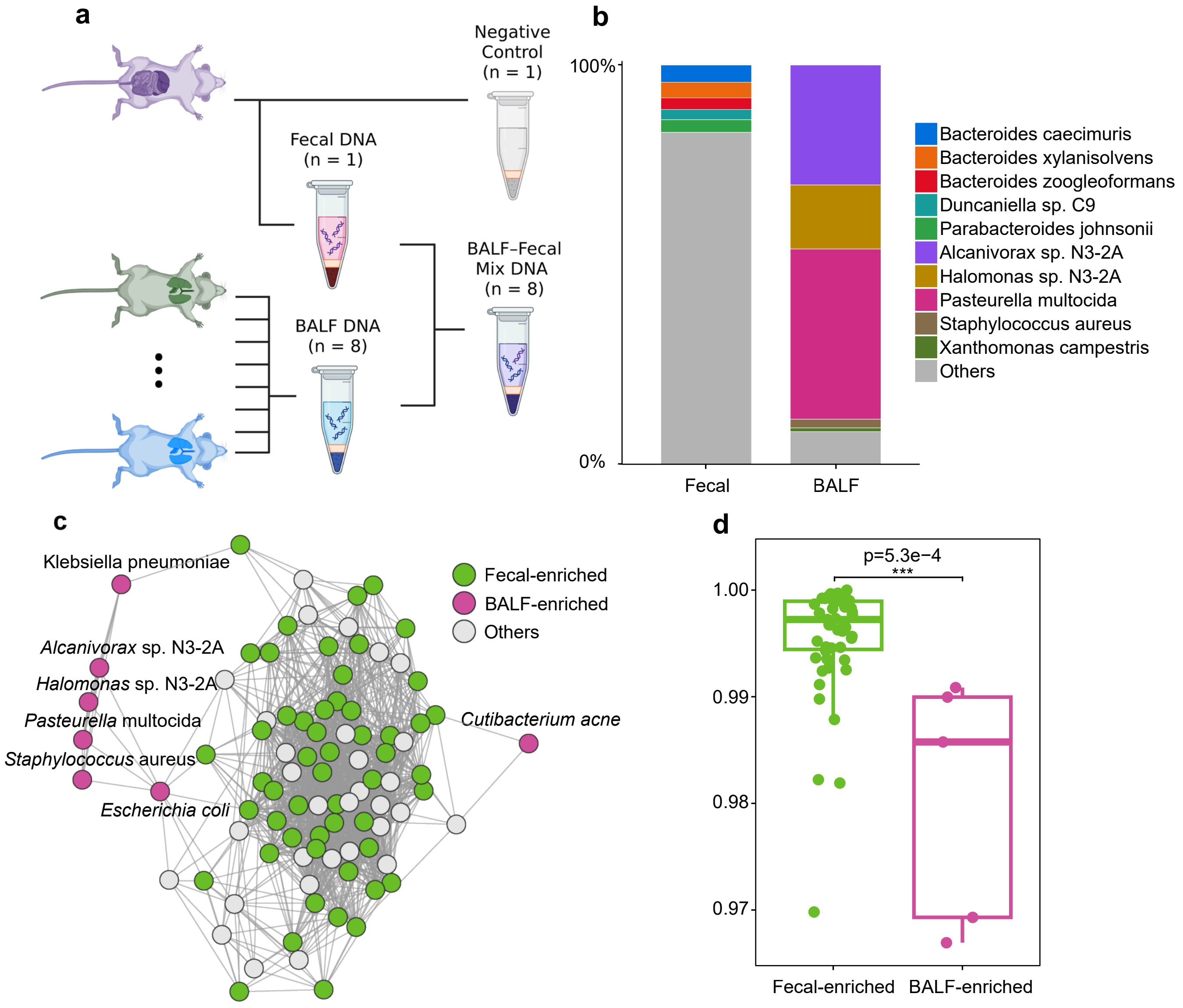


**Supplementary Fig. 9 | Experimental validation of Metacontam in a mouse BALF–fecal mixing study.**
**a,** Experimental design schematic for the mouse BALF–fecal mixing study. BALF DNA was extracted from eight mice (M1–M8) and each BALF DNA was mixed with fecal DNA from mouse M8 to generate eight mixture samples. Three additional controls were sequenced: fecal kit negative control, M8 fecal-only and M6 BALF-only. **b,** Stacked bar plot showing relative abundance of the top five species in fecal-only and BALF-only samples. For fecal-only, taxa detected in the kit negative control were excluded prior to ranking; remaining taxa are grouped as “Other.” **c,** Microbial association network inferred from the eight mixture samples. Species enriched in fecal-only or BALF-only samples are highlighted as fecal enriched or BALF enriched, respectively; all other taxa are shown in grey. **d,** ANI distributions for fecal-enriched and BALF-enriched species highlighted in c. Each point represents the per-species mean ANI across samples. Statistical significance was assessed using a one-sided Wilcoxon rank-sum test.

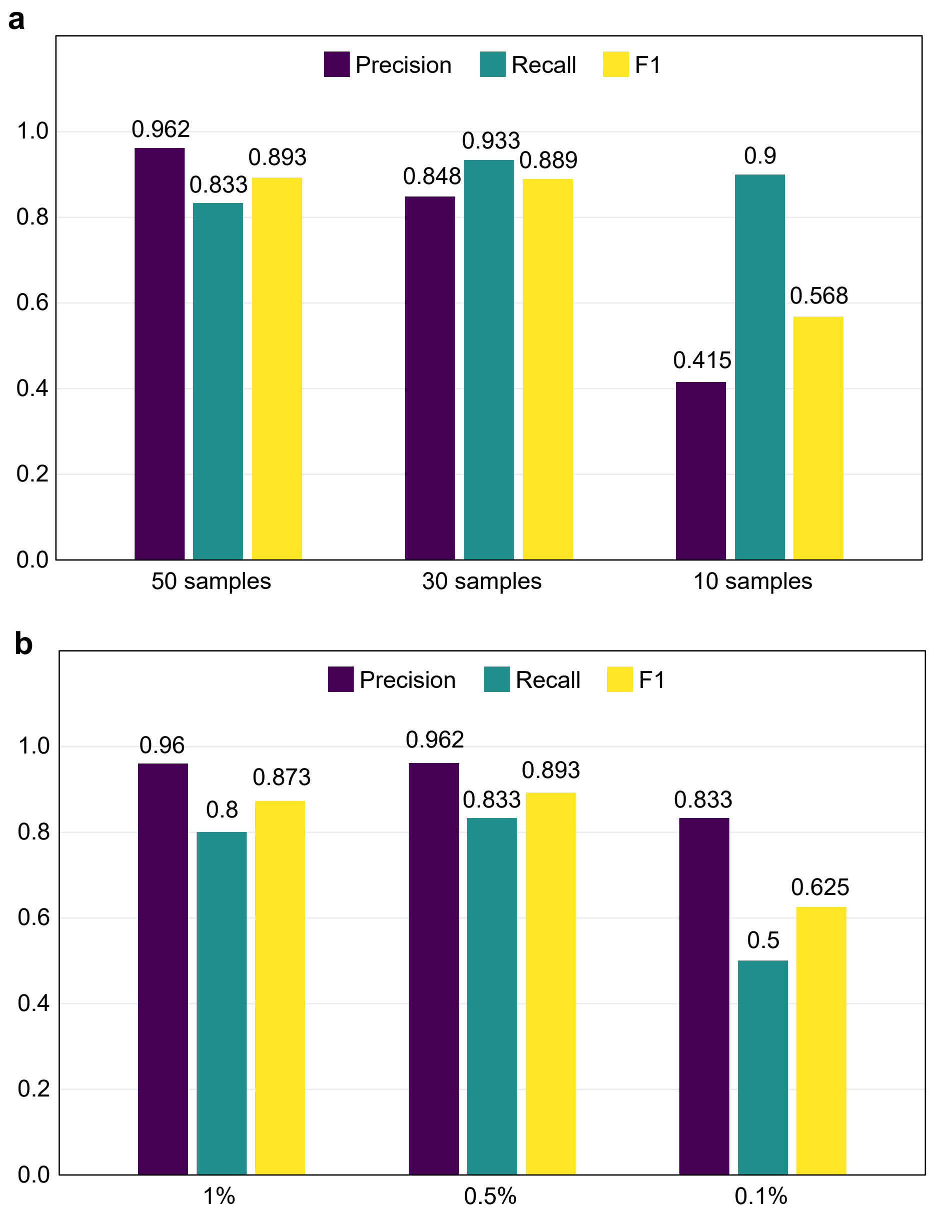


**Supplementary Fig. 10 | Simulation performance across sample sizes and spike-in levels.** Simulated fecal shotgun metagenomes were generated by spiking reads from 30 blacklist species into native fecal read sets (see Methods). **a,** Effect of sample size at fixed spike-in fraction (0.5%). Bars show precision, recall and F1 score against the ground-truth set (30 species) for n = 10, 30 and 50 samples. **b,** Effect of spike-in fraction at fixed sample size (n = 50). Bars show precision, recall and F1 score for spike-in fractions of 1%, 0.5%, and 0.1%.
